# Sickness behaviour within cleaning interactions

**DOI:** 10.1101/2024.04.17.589989

**Authors:** Tânia Marquês, Beatriz Pereira, Mélanie Marques, Tiago Repolho, José Ricardo Paula

**Author notes:** Correspondence: T. Marquês, J. R. Paula.

## Abstract

When facing diseases, strategies such as social distancing have proven effective in preventing disease spread. Cleaning interactions are social interactions where cleaner organisms remove ectoparasites from clients. These interactions can be potential vectors of infectious diseases in reef fishes. Here, we aimed to determine if cleaner wrasses (*Labroides dimidiatus*) employ social distancing and other behavioural tactics to prevent disease spread within the context of cleaning interactions. To this end, we used bacterial lipopolysaccharides (LPS) immunostimulation to induce “sickness” behaviour in cleaners. After LPS injection, we performed four behavioural trials: social preference (from cleaner and client perspectives), social interaction, and bystander cooperativeness tests. In the social preference test, LPS-injected cleaners showed lower activity rates but no change in their social preference. However, some clients preferred “sick” cleaners. Contrarily, during the social interaction test, cleaners reduced the number of interactions, despite clients’ willingness to interact. For the bystander test, LPS-injected cleaners were unable to adjust their behaviour to be more cooperative. Reduced activity levels revealed lethargy, which may lead to passive self-isolation, allowing for some social distancing of healthy individuals. This suggests that immune-stimulated cleaners undergo changes in their behaviour which can be viewed as a passive social distancing preventing disease spread.

## Introduction

In an unprecedented way, the recent COVID pandemic brought to our attention the need to consider the costs of social living, such as disease transmission, and, similarly, what measures social species can use to prevent these costs [1, 2]. Considering how social interactions create opportunities for diseases to spread, strategies can be employed to prevent disease transmission, such as early recognition of infected individuals and associated measures that reduce the costs of infections [3, 1]. One possible tool for early disease recognition is the identification of common sickness behaviours exhibited by infected individuals. These behaviours encompass several modifications including lethargy, somnolence, anorexia, as well as withdrawal from social interactions, characterized by a drop in interest in social activities [3, 4, 5].

Several factors may facilitate disease spread in the context of social interactions. Reduced spatial distance and the specific locations where interactions occur, such as doctors’ waiting rooms, where there is a higher infection pressure, increasing the risk of transmission for those who attend such disease hotspots [6, 7, 5, 1].

Social distancing is a basic, low-cost behavioural strategy that individuals can use to mitigate disease spread [8, 1, 2]. Nevertheless, contrary to initial perceptions, humans are far from being the only animal that increase the distance between individuals (social distancing) and employ active behavioural strategies to avoid disease [5, 9, 2]. Sickness behaviour, active self-isolation, social avoidance and exclusion of sick individuals, and other adaptative strategies, including changes in appearance, smell and vocalisations, are commonly used [5, 9]. This shows how looking at disease avoidance behaviours in non-human animals is important, as they can provide insights into ecological and evolutionary processes between sociality and disease transmission [9]. It is also important to consider the costs of social distancing and how living in groups can leverage even when there is a high risk of infection. While social isolation can lead to increased stress, group living allows, among other benefits, for more effective predator vigilance and efficient foraging, justifying that social distancing could be disregarded [10, 2].

Cleaning interactions involve a cooperative exchange where an organism—the cleaner—removes ectoparasites, bacteria, and diseased or injured tissue from a host—the client. These interactions benefit both partners; the first uses the interaction as a food source while the second improves its health. Because of the benefits these interactions bring, they may be interpreted as examples of socially relevant interactions [11]. Furthermore, cleaning mutualisms represent a key role in coral reef ecosystems by reducing ectoparasites’ abundance and infestation rates, which translates into improving client fish fitness, experimental removal of cleaners from the wild showed a direct impact on client fish, resulting in slower growth rates and decreased abundance and diversity [12, 13, 14, 15].

Cleaners have the potential to spread diseases, as evidenced by certain wrasse species used as cleaners in fish farms [11]. Since the likelihood of parasite transmission is directly related to the number of daily contacts, the bluestreak cleaner wrasse, *Labroides dimidiatus*, which engages in a high number of daily interactions, has been considered to be a potential carrier of parasites to new clients, a phenomenon known as the “cleaners as transmitters” hypothesis described by Narvaez et al [16]. Furthermore, the parasite hotspot hypothesis suggests that parasite infection pressure may be higher in areas near cleaning stations, which can represent disease hotspots [16]. This way, the frequency and duration of cleaning interactions must be balanced so that cleaners don’t act as “superspreaders”, akin to healthcare providers, who also face the risk of transmitting pathogens to those they interact with [9].

In this study we aimed to investigate whether cleaner wrasses (*Labroides dimidiatus*) use social distancing and other behavioural tactics to prevent the spread of disease during cleaning interactions. To achieve this, we used bacterial lipopolysaccharides (LPS) immunostimulation to induce sickness behaviour in cleaner wrasses without causing harm to the fish. After the LPS immunostimulation, we conducted four behavioural trials: the social preference test (from both cleaner and client perspectives), the social interaction test, and the bystander cooperativeness test.

## Material and Methods

### Acclimation

All fish species (cleaner wrasse *Labroides dimidiatus*, n=23; clients *Acanthurus leucosternon*, n=3, *Naso elegans*, n=8, *Zebra*soma *scopas*, n=9, *Dascyllus trimaculatus*, n=5) were captured in Maldives and obtained through a commercial supplier, being transported under controlled conditions (temperature, dissolved oxygen, salinity) to our aquatic facilities. Upon arrival, cleaner wrasses *Labroides dimidiatus* were kept in individual tanks and provided with a PVC (polyvinyl chloride) tube for refuge (approximately 2 cm diameter and 5 cm length). The clients were laboratory-acclimated in common tanks. All tanks were maintained in seawater conditions similar to the collection site: salinity = 35±0.5, temperature 26°C, pH 8.1 and photoperiod of 12h/12h. Natural seawater (NSW) was pumped from the sea into a 5 m^3^ storage tank, filtered (0.35 μm) and UV-irradiated (Vecton 300, TMC, Portugal). Temperature was regulated using submerged seawater heaters (300W, TMC, Portugal) in temperature-controlled rooms. Additionally, the seawater abiotic conditions were monitored manually on a daily basis: temperature and oxygen (VWR® DO220, VWR, Portugal), salinity (HI98319 salinity tester, Hanna Instruments, Portugal) and pH (pH1100H, VWR, Portugal). Following three days of acclimation, cleaners were trained to feed on plexiglass plates.

### Immune stimulation

For the first experimental stage, on the third day of experiments and following a two-day learning phase of the bystander test (see below), cleaners were randomly divided into two groups of 15 individuals each (the remaining individual was used in intraspecific tests, check below). One group received treatment with LPS, while the other served as the control treatment and received phosphate-buffered saline. LPS or saline was administered via intraperitoneal injections using heparinised 29-gauge insulin syringes (heparin concentration: 28 mg ml^-1^). For the LPS treatment fish, a dosage of 30 mg kg^-1^ was used (Sigma-Aldrich L2880, serotype 055: B5), while control fish were injected with a 0.01 µmol PBS (Sigma-Aldrich, Steinheim, Germany), following the protocol outlined by Binning et al. [17]. For both treatments, the injection volume was mass-adjusted for each fish [17, 18].

After performing the experiments, the spleen-somatic index (SSI) was used to measure immune activation in the fish. Cleaner fish were euthanised with an overdose of MS-222 (morphine sulfate) (Acros Organics, Geel, Belgium, at a concentration of 0,5 g per 200 ml). Total body and spleen weights were measured using a precision scale (Shimadzu, AUW220D). Total length (TL), and standard length (SL) were measured using a calliper (Vernier Analog Calliper 150 mm, Macfer). The spleen-somatic index was calculated using the following formula [17, 19, 20].

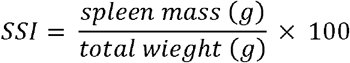

### Bystander test

Since previous experiments demonstrated how cleaner wrasses are more cooperative in the presence of an audience [21], this experimental test assessed whether “sick” cleaners could adjust their feeding behaviour when an image-scoring bystander was present. Plexiglas plates were used to simulate the presence of other reef fish as clients, offering two types of food items: prawn (which represented the preferred food item equivalent to the client’s mucus and, when chosen, is considered cheating) and a mix of fish flakes and prawn (hereafter referred to as “flakes”, representing less preferred ectoparasites) [15, 22].

The “bystander test” included two phases: a learning phase and a test phase. For the learning phase, there were six trials in total (two trials on day one and four on the second day of the experiment), with each trial being separated by at least one hour. In each learning trial, a Plexiglas plate with fourteen food items (twelve flakes and two prawns) was presented to the individual cleaner. If the cleaner fed on prawn the plate was removed for 60 seconds before being made available again. If the cleaner cheated a second time, the plate was removed until the next trial. However, if the cleaner only fed on flakes, the plate would remain available without being removed. The purpose of the learning phase was to train the cleaner to feed on flakes instead of prawns [22].

The test phase is intended to determine if cleaners can adjust their feeding behaviour when an image-scoring bystander is present. During the test phase, which started on day four of the first experimental stage, cleaners were presented with new plates similar to those used in the training phase. However, this time, half the plates had a yellow and green stripe, while the other half had a grey stripe, each containing four food items, two flakes, and two prawns. The test phase involved two scenarios: a single plate scenario (control) where the cleaners were presented with a single plate representing the client, and a double plate scenario (treatment) where they were exposed to two plates simultaneously, with one representing the client and the other representing the bystander (an “image-scoring situation”). In both cases, the plate(s) were removed immediately if a prawn item was consumed and remained if flakes were eaten. In the image-scoring scenario, cleaners had access to a bystander plate to increase their image score and demonstrate cooperation. Cleaners could achieve this by only feeding on flake items on the first plate and then having access to the second plate, eating four flake items before eating the prawn item. The test phase consisted of five rounds, each with two trials (the single plate and the double plate scenario) spaced 30 minutes apart, with the following round starting 60 minutes later. Four rounds were performed on day four, and the fifth on day five. The colour of the stripe (yellow and green/grey) for the “image-scoring situation” and the position of the plate (left/right) were randomly assigned. At the beginning of each trial, the subjects were confined to the back of the aquarium using an opaque partition that was removed once the plates were in place [22].

The learning phase was conducted on the days immediately before the day of the injection with LPS, and the test phase trials started the day after the injection (four trials on the first day and the last one the following day). The bystander effect score was calculated by subtracting the control trial ratio (i.e., number of flake items divided by the number of prawn items eaten) from the treatment trial ratio (i.e., number of flake items divided by the number of prawn items eaten) [22].

### Social preference

The social preference test is intended to determine an individual’s preference for social stimuli. It involved two phases: a habituation phase, during which an individual was placed in a tank to explore an environment devoid of any social stimulus, and an interaction phase, during which a social stimulus was presented to the individual.

In this study, cleaners and clients were submitted to the social preference test. For the cleaners, two different experimental treatments were used: one tested the preference between a client (*Acanthurus leucosternon*) and a novel non-social object (NO), and the other tested the preference between a conspecific and a NO [23, 24]. To perform the test, rectangular-shaped tanks were used (81 cm length x 18.5 cm high x 20.5 cm wide), featuring two smaller lateral chambers (15.5 cm) and a larger central chamber divided by transparent partitions to allow visual contact with the stimuli. The central chamber was marked with tape to indicate the area near both stimuli. After the stimuli were in place, an opaque partition was used to hide them. The individuals were placed in the central chamber for the habituation phase (5 minutes), after which the opaque partition was removed. The interaction phase began, and the individuals were free to move within the chamber for 10 min, recorded by a SONY camera (HDR-CX240E, 9.2 Megapixels) from above. After completing both treatments, the seawater of the central chamber was replaced before another individual was tested, with the stimuli alternating location among rounds. These tests were conducted on days five and six of the experiment.

For the clients (*Dascyllus trimaculatus*, n=5, *Naso elegans* n=8 and *Zebra*soma *scopas*, n=9), social preference was tested in a second experimental phase, following the same procedure as cleaners. The clients were presented with two cleaners (i.e., as social stimuli), one injected with LPS and the other with saline (control), with interactions also recorded from above, but using a GoPro Hero 3+.

Social preference was inferred by measuring the percentage of the total time the individual spent in proximity to the stimuli. Videos were analysed using AnimalTA, which allowed several variables to be obtained, such as the total duration of the test, time spent near the stimuli and in the central area, and cleaners’ activity levels, measured by the total distances roamed [25].

### Interaction

The interaction test was designed to assess the interaction between an individual (cleaner) that has been immune-stimulated (i.e. has an activated immune system) and a fish species that is commonly serviced by this individual in their natural environment (i.e. the “client” fish, namely the *Acanthurus leucosternon*), by placing them together in a tank. This test began with a habituation phase (5 minutes), during which both fish were separated by an opaque partition to acclimate to the environment. After the partition was removed, the interaction phase began, lasting a minimum of 15 minutes. During this time, the behaviour of both fish was recorded (SONY, HDR-CX240E, 9.2 Megapixels; CANON EOS 350D, 8.0 Megapixels) and later analysed. Interaction tests occurred following the cleaners’ social preference test on days six and seven of the experiments.

The recorded videos of the interactions were analysed using Boris software [26], facilitating the identification of different behaviours, namely i) chases (when the client fish chases the cleaner as a form of punishment after it cheats), ii) interactions started by the cleaner (when the cleaner approaches the client and initiates the interaction), iii) interactions started by the client (when the client approaches the cleaner and initiates the interaction), iv) client jolts (sudden movements of the client in response to something the cleaner did, such as removing mucus), v) posing (when the client positions itself submissively to demonstrates willingness to be cleaned), vi) cleaning (when the cleaner removes parasites from the client), vii) tactile stimulation (when the cleaner uses its pectoral fins to stimulate the client), and viii) dance (the oscillatory swimming movements of the cleaner to demonstrate availability to clean) [27].

### Statistical analysis

To perform data exploration, and following Zuur & Ieno [28], generalised linear models (GLM) were used to analyse behavioural data, namely SSI, Bystander-effect, Total Distances and Interaction variables (number of interactions, proportion of interactions started by cleaners, proportion of client jolt events per second, proportion of posing displays by clients) with LPS injection treatment (factor with two levels: injected and control), using Gaussian distribution for these variables (excluding number of interactions where Poisson distribution was used). For all Social Preference scenarios, and to analyse the time spent in the proximity of the stimuli, the function ‘glmmTMB’ from the package ‘glmmTMB’ was used [29], and Post hoc multiple comparisons were performed using the package ‘emmeans’ [30] with Tukey corrections.

Except for the Bystander-effect, for which it was used the function “T-Test”, the treatment effects for all variables where analysed using the function “Anova” [31]. Model assumptions and performance validation used the package “performance” [32], with the analysis being performed in R, version 3.4.3 (R Core Team, 2017) and using the HighstatLibV10 R library from Highland Statistics [33].

## Results

### Immune response

Cleaner fish injected with LPS had a significantly higher SSI than saline-injected fish (*χ*^2^ = 7.060, d.f. = 1, p = 0.008, Fig. S1).

### Bystander test

Regarding the bystander test, saline-injected cleaners exhibited, on average, higher bystander scores than LPS-injected cleaners. Here, solving the test implies having an average score higher than zero, results show that while the control group had a score significantly different from zero (t = 2.333, d.f. = 9, p = 0.044, mean = 0.280), while LPS-injected did not differ from zero (t = -0.238, d.f. = 6, p = 0.8194, mean = -0.031) (Fig.1).

**Figure 1.**
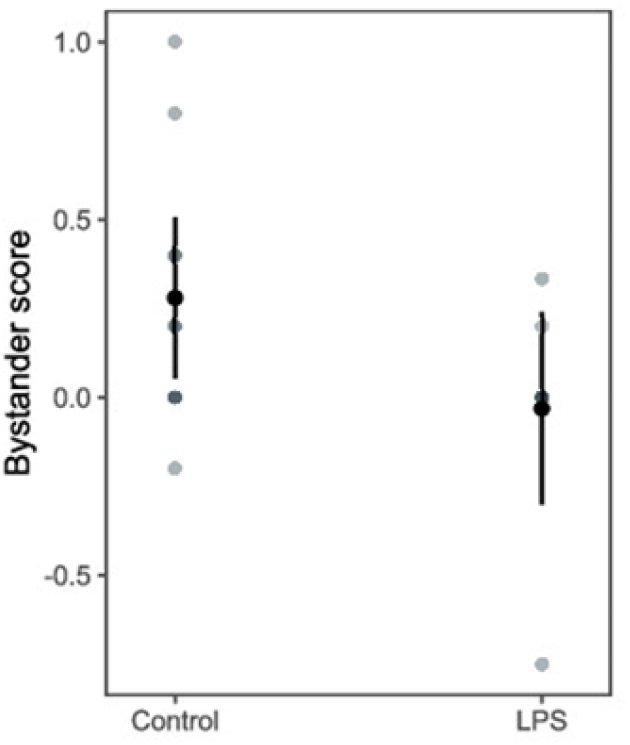
Bystander score for control and LPS-injected individuals. Back-transformed predicted means ±95% C.I. from the model and raw data values are presented.

### Social Preference

In social preference test performed on cleaners, cleaners injected with LPS swam shorter distances than cleaners in the control group, regardless of the social stimuli presented to them (client: *χ*^2^ = 5.192, d.f. = 1, p = 0.023, Fig. 2A; cleaner: *χ*^2^ = 7.648, d.f. = 1, p = 0.006, Fig. 2B).

**Figure 2.**
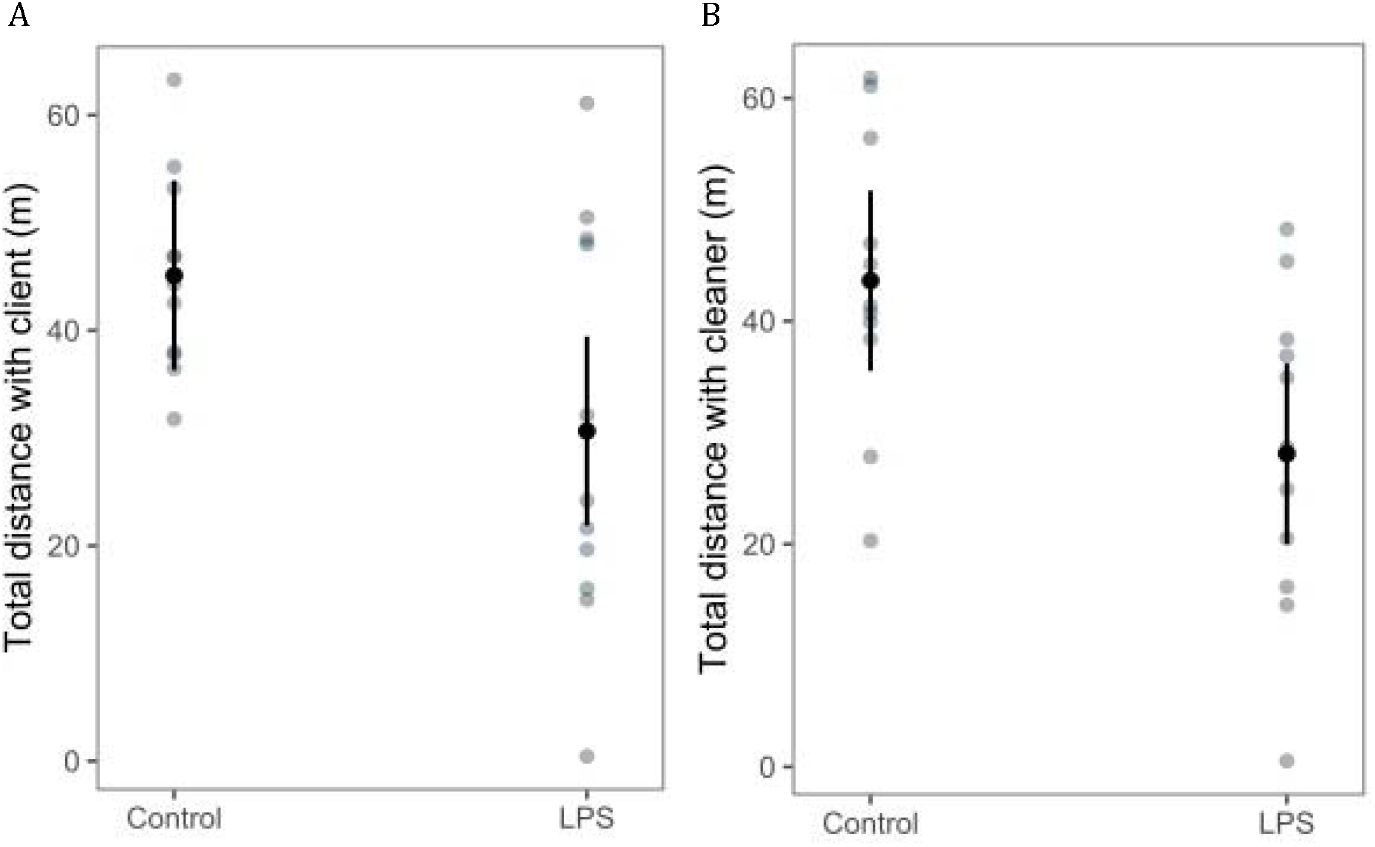
Social preference of control and LPS injected cleaners, in the scenarios of client (*Acanthurus leucosternon*) versus NO and of conspecific (other cleaner) with NO. A) total distance, in meters (m), roamed when with client; B) total distance, in meters (m), roamed when with other cleaner. Back-transformed predicted means ±95% C.I. from the model and raw data values are presented.

Regarding the time spent near the stimuli, cleaner fish spent more time in the client’s presence (*Acanthurus leucosternon*) than in the presence of the NO, regardless of the treatment cleaners had been submitted to (control or LPS). However, in the scenario of conspecific *versus* NO, there were no significant differences in the time spent near either stimulus. This was despite the difference in time spent by control individuals in the proximity of a conspecific or the neutral area (Table 1, Table 2, Fig. 3).

**Table 1.**
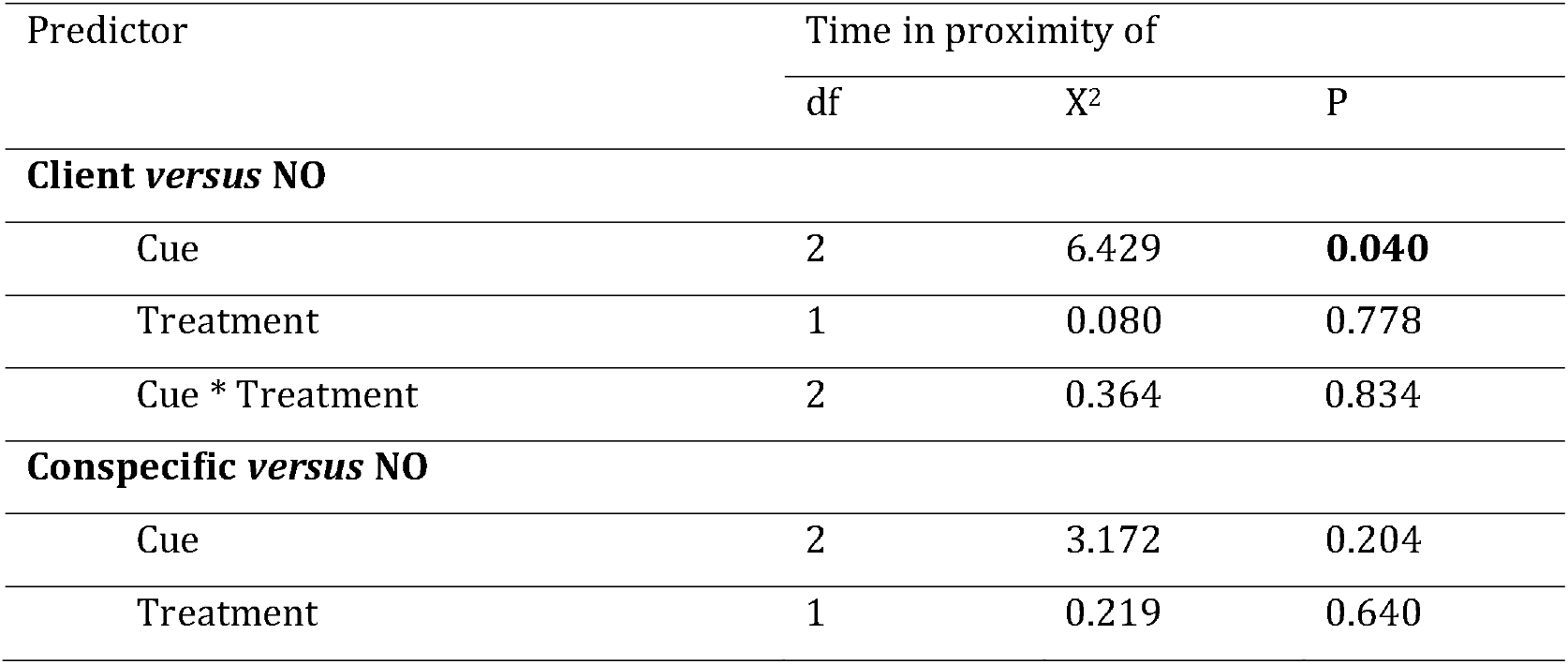
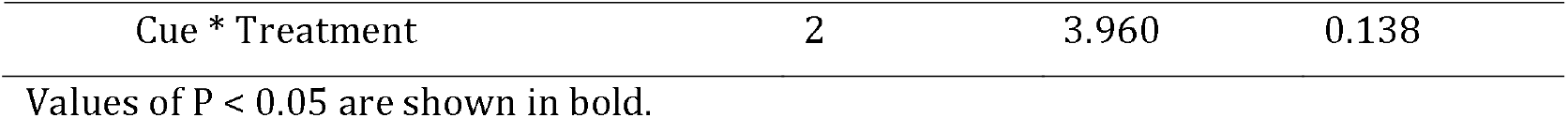
Analysis of deviance table (type II tests) for cleaners’ social preference.

**Table 2.**
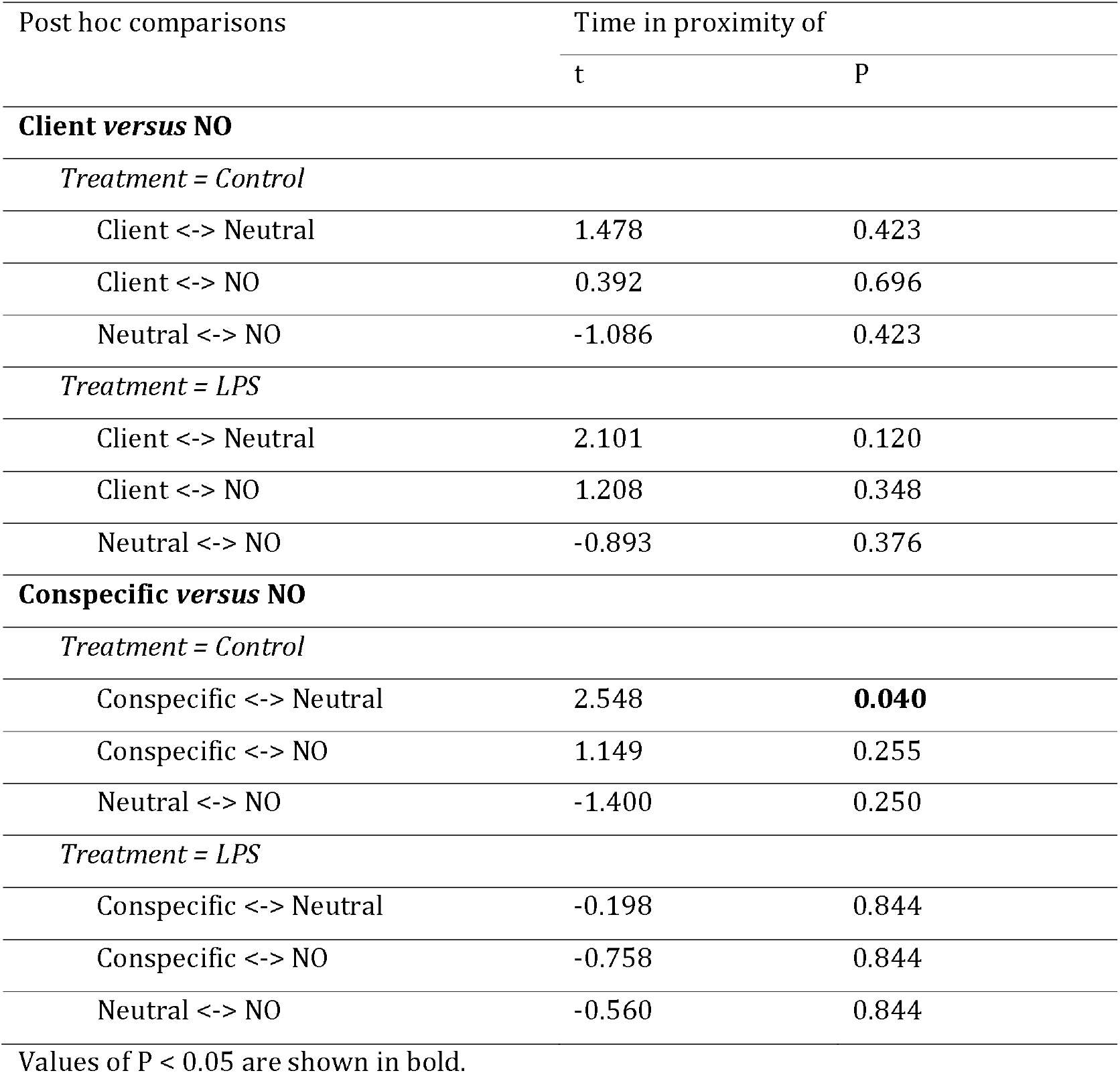
Contrasts of the post hoc multiple comparisons for cleaners’ social preference.

**Figure 3.**
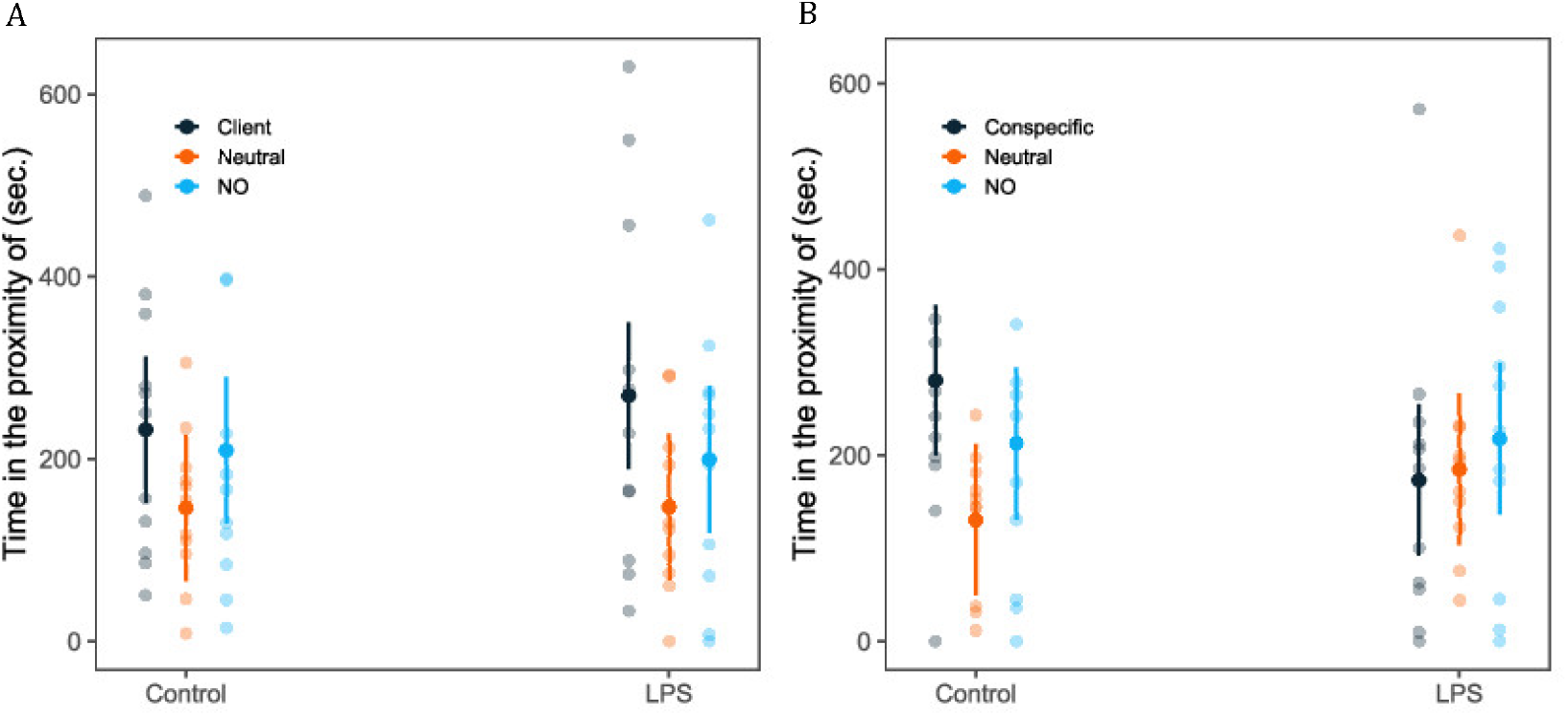
Social preference of control and LPS injected cleaners, in the scenarios of the client (*Acanthurus leucosternon*) *versus* NO and conspecific (other cleaner) *versus* NO. A) time, in seconds (sec.), spent near each stimulus (Client, Neutral and NO); B) time, in seconds (sec.), spent near each stimulus (Conspecific, Neutral and NO); Back-transformed predicted means ±95% C.I. from the model and raw data values are presented.

In the assessment of clients’ social preference, variations in the time spent near each stimulus (control or LPS-injected cleaners) were observed, regardless of clients’ species. Specifically, the time spent by *Dascyllus trimaculatus* near LPS-injected cleaners or in the neutral area, as well as in time spent by *Zebrassoma scopes* near control and LPS cleaners and near LPS cleaners and in the neutral area (Table 3, Table 4, Fig. 4).

**Table 3.** Analysis of deviance table (type II tests) for clients’ social preference.

| Predictor | Time in proximity of |  |  |
| --- | --- | --- | --- |
|  | df | X<sup>2</sup> | P |
| Cue | 2 | 11.741 | **0.002** |
| Species | 2 | 0.083 | 0.959 |
| Cue * Species | 4 | 8.167 | 0.086 |

Values of P < 0.05 are shown in bold.

**Table 4.** Contrasts of the post hoc multiple comparisons for clients’ social preference.

| Post hoc comparisons | Time in proximity of |  |
| --- | --- | --- |
|  | t | P |
| Species = *Dascyllus trimaculatus* |  |  |
| Control <-> LPS | -2.135 | 0.056 |
| Control <-> Neutral | 1.093 | 0.280 |
| LPS <-> Neutral | 3.228 | **0.006** |
| Species = *Naso elegans* |  |  |
| Control <-> LPS | -0.675 | 0.754 |
| Control <-> Neutral | -0.975 | 0.754 |
| LPS <-> Neutral | -0.300 | 0.765 |
| Species = *Zebrassoma scopas* |  |  |
| Control <-> LPS | -2.367 | **0.032** |
| Control <-> Neutral | 0.194 | 0.847 |
| LPS <-> Neutral | 2.561 | **0.032** |

Values of P < 0.05 are shown in bold.

**Figure 4.**
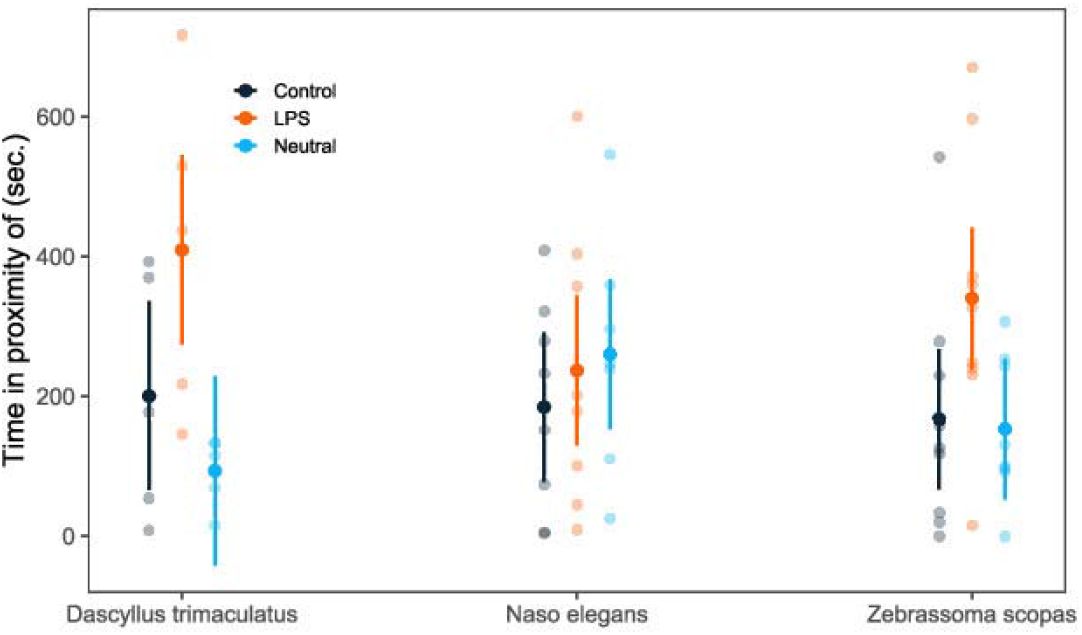
Social preference of clients’ species (*Dascyllus trimaculatus, Naso elegans and Zebrasoma scopas*). Time, in seconds (sec.), spent near each stimulus (Control = saline injected cleaner, LPS = LPS injected cleaner and Neutral area). Back-transformed predicted means ±95% C.I. from the model, and raw data values are presented.

### Interactions

LPS injection significantly lowered the number of interactions (*χ*^2^ = 14.206, d.f. = 1, p = 0.0002, Fig. 5A), with the proportion of client jolts events per second also being significantly altered by LPS injection (*χ*^2^ = 4.958, d.f. = 1, p = 0.026, Fig. 5C). However, no significant differences were found between control and LPS-injected cleaners regarding the proportion of interactions initiated by cleaners (*χ*^2^ = 1.324, d.f. = 1, p = 0.250, Fig. 5B), or the proportion of posing displays by clients (*χ*^2^ = 1.142, d.f. = 1, p = 0.285, Fig. 5D).

**Figure 5.**
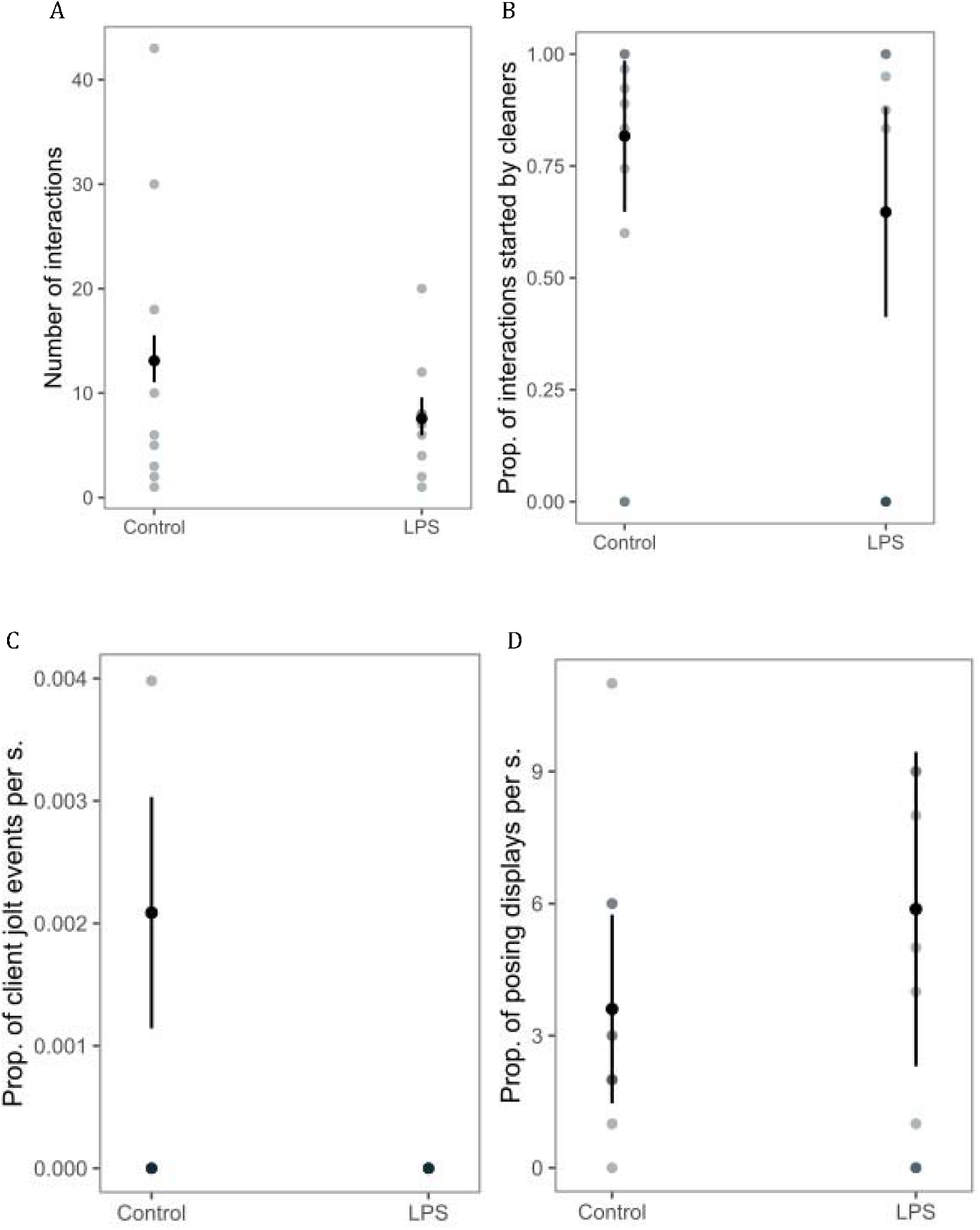
Behavioural responses observed during the interaction test between the cleaner (*Labroides dimidiatus*) and the client (*Acanthurus leucosternon*) A) number of interactions; B) proportion of interactions initiated by the cleaner; C) proportion of client jolt events per second; D) proportion of posing displays by clients. Back-transformed predicted means ±95% C.I. from the model, and raw data values are presented.

## Discussion

The induction of sickness behaviour in cleaner wrasses aimed to determine whether active behavioural strategies are employed to mitigate disease spread within cleaning mutualisms. Following LPS injection, cleaner wrasses exhibited a higher SSI than saline-injected cleaners, indicating that LPS injection successfully induced immunostimulation. As anticipated, immunostimulation induced sickness behaviour in cleaners, with *L. dimidiatus* being less active (lower distances travelled, Fig. 2A and Fig. 2B), which suggests that pathogen-free LPS immunostimulation is a suitable method for studying sickness behaviour.

According to Stockmaier et al. [9], behavioural responses resulting from immune challenges possibly stem from a combination of factors, including modifications in social cues and infection signs. While social hosts, may disguise the infection condition to continue benefiting from group living, cleaners failed to conceal their sick status (Fig. 1 & Fig. 2). This could be attributed to the intensity of the symptoms induced by LPS immunostimulation [4, 10], or other factors such as gender or social rank [3]. Additionally, the interplay between immune system function and neurological disturbances may contribute to behavioural changes [24]. Furthermore, passive-self isolation due to indirect physiological responses to infection, such as lethargy, can modify movement patterns and reduce contact with other individuals [9]. When feeling sick, individuals tend to be less likely to engage in social relationships due to lethargy. This phenomenon is widely observed across species, highlighting the interplay between sickness behaviour and social context, as well as between immune response and exploratory and social behaviour [7, 24, 4, 9]. Therefore, the difference in activity levels of these immune-stimulated cleaners can be attributed to the lethargy characteristic of sickness behaviour.

In social preference tests, the behaviour of injected cleaners varied depending on the scenario. When faced with a choice between a client and a non-social object (NO), there were significant differences in the time spent in proximity to each stimulus. However, in the scenario of a conspecific versus a NO, the time spent in the proximity of the stimulus did not show variance (Table 1, Table 2, Fig. 3). These results underscore the natural tendency of cleaners to prioritise cleaning interactions with clients, regardless of their own health status. Cleaners are expected to maintain sociability and favour social interactions as highly social animals that engage in thousands of daily interactions [10]. Therefore, by abstaining from social distancing, cleaners prevented potential conflicts of interest that might arise from such behaviour [9]. By suppressing their sickness behaviour, individuals gain direct fitness benefits and sustain their normal functioning [3]. Moreover, the influence of social environment (e.g., the rich social life of complex societies where individuals are widely stimulated during their early development stages) must be considered. Such environments modulate behaviour, providing a background that encourages individuals to diminish or even forego sickness behaviour [8, 3, 34]. By considering the changes in physical, environmental, and social factors and how these will influence individual behaviour, group formation, and sociality, one can better evaluate the possibility of highly social individuals, such as cleaners, to include social distancing as part of their behavioural repertoire [35, 36, 37].

LPS-injected cleaners engaged in fewer cleaning interactions (Fig. 5A), potentially indicating social distancing or increased lethargy. The higher number of client jolts (i.e., dishonesty) observed for control group cleaners compared to LPS-injected ones (Fig. 5C), suggests that the reduction of interactions in LPS-injected cleaners may be related to lethargy. Additionally, decreased cooperativeness was also evident in the bystander test (Fig. 1), where LPS-injected cleaners failed to adjust their behaviour cooperatively, unlike control cleaners. Despite this, client fish maintained their intention to socially interact with LPS-injected cleaners (i.e. posing displays, Fig. 5D), suggesting a lack of measures on behalf of the client to prevent disease transmission, such as avoidance behaviours [9].

Previous interactions influence mutualism stability, with clients returning to the cleaning station where there was no conflict. Additionally, studies have shown that the quality of service provided by cleaners is influenced by their interaction with previous clients, with short-term stress often resulting in heightened levels of cooperation [38, 39]. In our study, although LPS-injected cleaners failed to adjust their behaviour towards the clients and manage their reputation effectively, clients continue to exhibit interest in engaging in cleaning interactions with these individuals. The extent to which these results demonstrate the inability of clients to detect sickness behaviour in cleaners or if they deliberately chose to interact with sick cleaners it remains uncertain.

Given that fish may be more susceptible to pathogen transmission during cleaning interactions, and considering the frequent, prolific, transient, and intimate nature of these interactions among cleaners, the lack of social distancing, and the absence of social avoidance among clients, cleaning interactions can indeed be viewed as potential ‘super-spreader’ events [40].

LPS-injected cleaners could not adjust their behaviour in the presence of an audience (Fig. 1), suggesting a potential decline in their highly sophisticated decision-making abilities [21, 41]. Similar cognitive impairments have been reported in other species affected by pathogens, highlighting the intricate relationship between behaviour and the immune system in fish [42, 24]. Moreover, the energy consumption associated with infection can lead to resource reallocation, further compromising cognitive function, which is further influenced by the severity of the infection [17]. Considering how cleaning mutualism relies on cognitive processes, it comes as no surprise that the finetuning of this interaction is negatively impacted by sick behaviour [15]. The extent to which the inability of cleaners to adjust their behaviour in the presence of a bystander is attributed to cognitive impairments alone or a combination of cognitive decline and socioecological pressures requires further investigation.

### Conclusions

In conclusion, despite the benefits of simple social adaptations like social distancing, which are known to reduce the costs of disease spread [9], there is no evidence of such measures in this cleaning mutualism. Furthermore, our findings suggest that clients failed to detect sickness behaviour, such as lethargy, and did not use it to control pathogens’ transmission [7, 9]. Moreover, the fact that LPS-injected cleaners did not adjust their behaviour in the presence of a bystander suggests that sickness may compromise cognitive sophistication and ability to manage their reputation [43].

Further studies could benefit from exploring the possible use of social distancing among cleaners by using immunostimulated cleaners as stimuli in the social preference tests. It would also be beneficial to understand how the behaviour of sickness in clients may influence the cleaning mutualism in both the social preference test and the interactions. Finally, further mechanisms studies are required to understand the neurobiological processes associated with sickness behaviour.

Our findings underscore the lack of social avoidance towards sick-behaving cleaners, reinforcing the characterisation of cleaning interactions as super-spreader events [40, 44].

