## Supplementary material for "Sickness behaviour within cleaning interactions": Fig. S1

### ***Supplemental Materials***

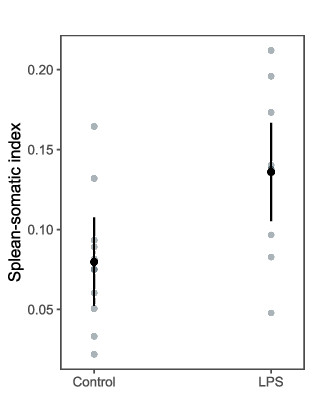

Figure 1 – Spleen-somatic index for control and LPS-injected individuals. Back-transformed predicted means ±95% confidence intervals (C.I.) from the model and raw data values are presented.
